# Using extracellular low frequency signals to improve the spike sorting of cerebellar complex spikes

**DOI:** 10.1101/556985

**Authors:** Gil Zur, Mati Joshua

## Abstract

The challenge of spike sorting has been addressed by numerous electrophysiological studies. These methods tend to focus on the information conveyed by the high frequencies, but ignore the potentially informative signals at lower frequencies. Activation of Purkinje cells in the cerebellum by input from the climbing fibers results in a large amplitude dendritic spike concurrent with a high frequency burst known as a complex spike. Due to the variability in the high frequency component of complex spikes, previous methods have struggled to sort these complex spikes in an accurate and reliable way. However, complex spikes have a prominent extracellular low frequency signal generated by the input from the climbing fibers. We exploited this to improve complex spike sorting by applying Principal Component Analysis (PCA) on the low frequencies of the signal and show that the low frequency first PC achieves a better separation of the complex spikes from noise. The low frequency data are more effective in detecting events entering into the analysis, and therefore can be harnessed to analyze the data with a larger signal to noise ratio. These two advantages make our method more effective for complex spike sorting. Our characterization of the dendritic low frequency components of complex spikes can be applied in other studies to gain insights into processing in the cerebellum.

## Introduction

One of the fundamental primary tasks applied on a dataset is to extract relevant events by separating the signal (event of interest) from noise. For neural activity in particular, the events of interest are often located in the spiking activity generated by the neurons. Thus, one of the key steps in analyzing neural data is to isolate the spike activity from the rest of the signal. A standard method for monitoring spike activity is to record extracellular activity. Spike sorting of extracellular activity can be quite challenging because of the multiple sources of variability. There are many extracellular spike shapes. Shape inconsistency can arise from non-stationary neural processes during burst or after a long pause (Elias et al., 2007). Shape inconsistency can also be the result of experimental measurement errors such as drifts in the location of the electrode (Gold et al., 2006). In addition to shape inconsistency, noise from other sources such as neighboring neurons or other electrical devices can corrupt the signal. Therefore, many analysis techniques and sorting methods have been developed to improve the quality of spike sorting analysis (Lewicki, 1998; Quiroga et al., 2004).

Nevertheless, the sorting methods used today only examine the high frequency components (typically > 300 Hz), since the low frequency signal is assumed to be uninformative. However, this assumption may not hold true; in particular for complex spikes of the Purkinje cells in the cerebellum that have low frequency components (Eccles et al., 1966; Warnaar et al., 2015). Complex spikes are driven by the input from the inferior olive which generates massive slow and large amplitude calcium spikes accompanied by bursts of high frequency spikes, known as spikelets. A second type of spike is called a simple spike. These spikes are modulated by the excitatory input from the parallel fibers (Voogd and Glickstein, 1998).

Generally, spikes are considered to be fast events occurring in a high frequency domain. The characteristic shape of the complex spike, together with presence of additional types of spike (i.e. simple spikes), make this signal challenging to detect using high frequencies data alone. The initial spike of the complex spike maybe similar to the characteristic shape of the simple spike. In addition, there is high variability in the number of spikelets and their timing (Yang and Lisberger, 2014), making this hard to exploit using an analysis that relies on shape features. The low rate of the complex spikes (~1 Hz) in comparison to simple spikes (~80 Hz) further challenges algorithms given the relative sparseness of the signal compared to the noise. Here, we present a novel approach to complex spike sorting based on the low frequency signal. Due to the highly elaborate dendritic tree of the Purkinje cell, the climbing fibers’ input generates an electrophysical signal composed of both low and high frequency domains. This low frequency signal often has a large, specific shape. We show how this signal can be exploited to improve complex spike detection and clustering from biological and non-biological noise. The findings show that in some cases low frequency signals can be used to substantially improve spike sorting.

## Methods

### Data acquisition

The data were collected from two male Macaca fascicularis monkeys (4-5 kg). All procedures were approved in advance by the *Institutional Animal Care and Use Committees* of The Hebrew University of Jerusalem, and were in strict compliance with the *National Institutes of Health Guide for the Care and Use of Laboratory Animals*. Monkeys were first trained to sit calmly in a primate chair (Crist instruments) and to consume food from a tube set in front of them. We implanted a headholder on the skull to allow us to restrain the monkeys’ head movements. In later surgery, we placed a recording cylinder stereotaxically over the floccular complex. The center of cylinder was placed above the skull targeted at 0 mm anterior and 11 mm lateral to the stereotaxic zero. We placed the cylinder with a backward angle of either 26 or 20 degrees (for each monkey).

Quartz-insulated tungsten electrodes were lowered into the floccular complex and neighboring areas to record simple and complex spikes using a Mini-Matrix System (Thomas Recording GmbH). Monkeys performed eye movement tasks during the recording (similar to Joshua and Lisberger, 2012). The analysis of the relationship between the responses of the cell and the behavioral task are reported elsewhere(Larry et al., 2019). Signals were high-pass filtered at the lowest frequency of 0.05Hz and digitized at a sampling rate of 40 kHz (OmniPlex). The dataset was composed of eight randomly selected Purkinje cells for which we manually inspected the data and marked the time point of each complex spike. These cells were used to test our sorting method. In addition, the data included 46 cells with well isolated complex spikes with a clear low frequency signal and a well isolated simple spike. These cells were sorted offline (Plexon); for sorting we used principal component analysis and corrected manually for errors. These cells were used to examine the properties of the complex spikes with low frequency signal.

### Analysis procedure-data filtering and setting a threshold

The first step of the analysis consisted of filtering the data with two separate high pass and low pass filters. For the high pass filter, we used a Butterworth filter with a cutoff frequency (−3 dB) of 600 Hz, and an order of 10. The low pass filter was the band-pass filter, which enabled us to remove very slow frequencies in addition to the high frequencies. The corner frequencies (−3 dB) for this filter were 20 Hz and 400 Hz, and the order of this filter was 2. The properties of the filter determine effectively spike sorting (Yael and Bar-Gad, 2017). Here we demonstrated an extreme case in which using the most common filter could remove almost all the information needed to detect the complex spikes. Thus, we verified that modifying the exact cutoff values used in the filters by changing the range of the corner frequencies (400-700 Hz for the high pass filter, and 200-500 for the low pass filter) did not alter any of the conclusions. In addition, the filtered data for both filters were compared to a finite impulse response filter (FIR) to verify similarity (high-pass with a corner frequency of 550 Hz, and an order of 1650; low-pass with corner frequencies of 30 Hz and 400 Hz, and an order of 12000). We refer the high pass filtered data as *high-frequency data*, and the low pass filtered data as *low-frequency data*.

After filtering the data, we set a threshold on the low-frequency data and looked for time points of threshold crossing. We then used these time points to select segments from the low frequency data around each threshold crossing time point. The segments were 10ms in length, and started 2ms prior to the threshold crossing. Using the same threshold crossing time points, we selected segments from the high-frequency data (starting 0.5ms prior to the threshold crossing, for a total length of 2ms), such that each segment in the low-frequency data had an equivalent segment in the high-frequency data. The high-frequency segments were aligned to the peak of the initial spike of the detected event. After the alignment, we updated the high-frequency segments such that each segment selected around its alignment point, within a window of 2ms-5ms, depending on the spikelets of the neuron. We categorized all the segments in two matrices: one for the low-frequency segments and the other for the high-frequency segments. We then performed principal component analysis (PCA) on each matrix.

### Analysis procedure-principal component analysis (PCA)

PCA is a common technique in spike sorting (Adamos et al., 2008). Its basic principle is to allow dimension reduction in order to visualize the data in a meaningful and useful manner. PCA reveals patterns in the data, by decomposing the data in a way that emphasizes the variation and distances within the dataset. The method makes it possible to find the direction with the largest variance to obtain a representation of the data in a reduced dimension that preserves the largest fraction of the variance (the distances within the dataset). The direction of the largest variance in the data is known as the first principal component (first PC). In the same way, the perpendicular direction to the first PC which has the largest variance is called the second principal component (second PC). We projected the data and looked for its coordinates in these directions.

The PCA on both datasets (low-frequency and high-frequency) identified the first PC for each dataset, and the coordinates of the datasets in these PCs. Recall that for each detected event (i.e. threshold crossing), two segments were selected: the low-frequency segment and the high-frequency segment. For each event we projected its low-frequency segment on the low-frequency’s first PC, and its equivalent high-frequency segment on the high-frequency’s first PC, to obtain the event score (coordinates) for these directions. For each event we then plotted the score on the low-frequency’s first PC against the score on the high-frequency’s first PC. This yielded a 2D representation of the data, where each dot represents a single threshold crossing event. This presentation included all the event of interest, and often separated data points into distinct signal and noise clusters. A polygon drawn manually around the 2D representation used to marks the complex spike cluster. To define the boundary of the polygon we examined the high and low frequency waveforms of dots around the margins of the polygon. This resulted in a better classification of the data, and made it possible to choose the clusters that contained the signal.

We developed a program for complex spikes analysis using low frequency signal which is available online (https://github.com/MatiJlab/ComplexSpikeDetection).

## Results

### Using separate low and high frequency components to sort complex spikes

We recorded extracellularly from Purkinje cells in the flocculus complex, and neighboring areas while the monkeys performed eye movement tasks (similar to Joshua and Lisberger, 2012). The complex spike extracellular signature contains both low and high frequency components (Warnaar et al., 2015). Figure 1 illustrates these components in two example neurons. We filtered the broadband extracellular signal (Fig. 1A and D) into low and high frequency components (Fig. 1B and E, 30*Hz* < frequency < 400*Hz* and Fig. 1 C and F, frequency > 600*Hz*). The low pass filter preserves the characteristic envelope shape of the complex spikes. The high pass filter captures the fast fluctuating events; i.e., the first large amplitude spike and the following spikelets. The two example neurons in Figure 1 had a clear low frequency component (Fig. 1B and E). If these events are indeed distinct from other extracellular events, they could be useful for identifying complex spikes. Thus, we examined whether we can use the low frequency components for detecting and sorting complex spikes.

**Figure 1:**
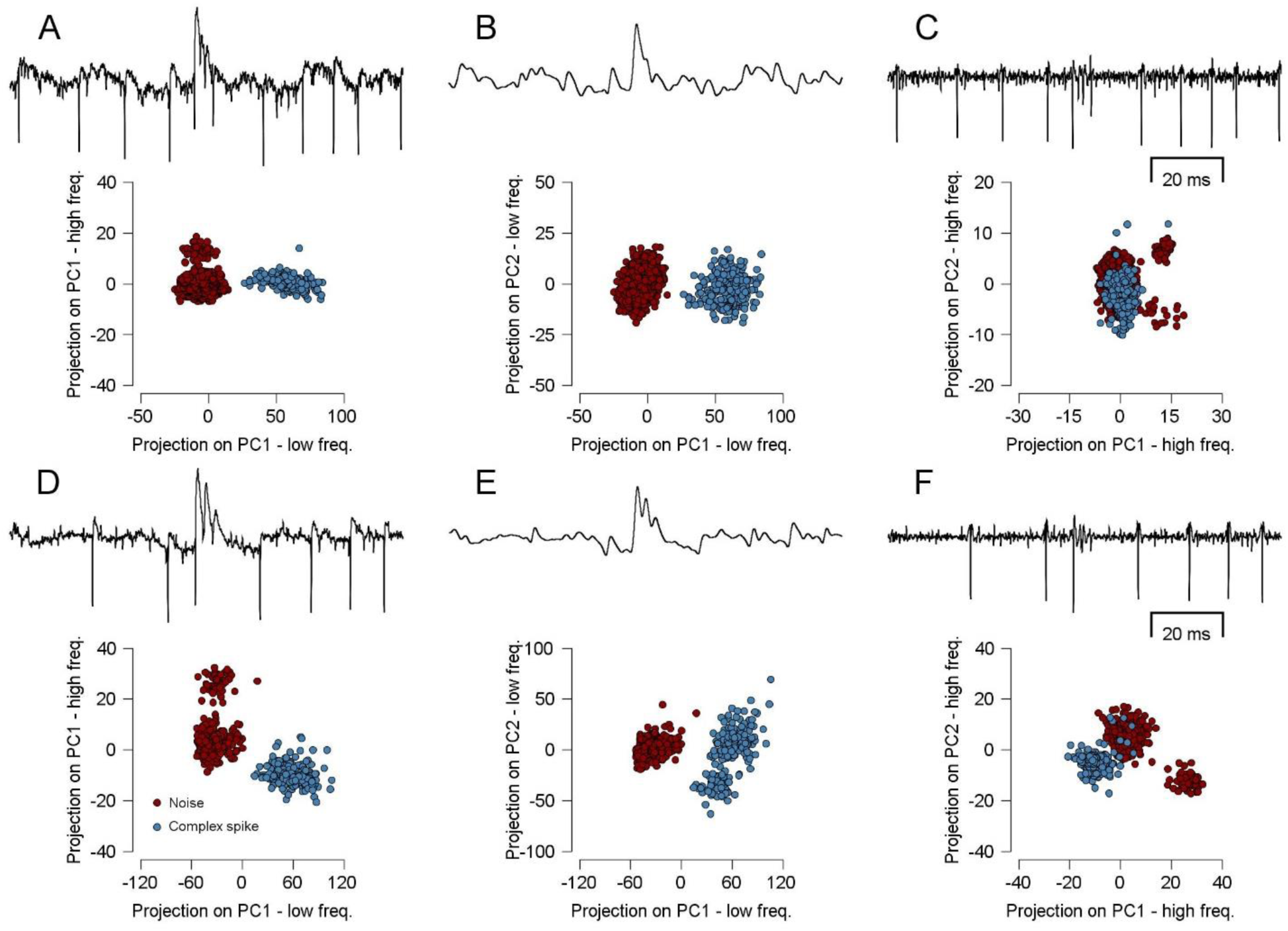
Illustration of signal and noise separation in several PCAs. Each row represents data from one neuron in three forms of PCA: (**A, D**) projection on low frequency 1^st^ PC and high frequency 1^st^ PC. (**B, E**) Projection on low frequency 1^st^ and 2^nd^ PC. (**C, F**) Projection on high frequency 1^st^ and 2^nd^ PC. Each dots represents a single complex spike (light blue) or noise events (red). In each row, a sample complex spike appears at the top of each scatter taken from the domain of frequencies used in the projection below it.

To sort the complex spikes, we extended Principal Components Analysis (PCA, see methods) (Abeles and Goldstein, 1977; Lewicki, 1998) to low frequency data. We performed PCA separately for the low and high frequencies and projected the data on the PCs to obtain the score (i.e., the coordinate) of each segment in the PC direction. We used either the low frequency (Fig. 1B, E) or high frequency (Fig. 1C, F) projections to sort the data. Selecting the leading PC separately for the low and high components reduces the initial dimensionally of the data to a common dimensionality. Thus, we can also use PCA to combine the projections on low and high frequency PCs (Fig. 1A, B).

The low frequency allows for a better separation of complex spikes from the noise. To illustrate the intuitive superiority of our analysis we compared different PCA methods in two example neurons (Fig. 1). We performed PCA and plotted the projections of the events on the PCs. We manually verified for each cluster whether it contained complex spikes or noise. To determine whether the PC projection separated complex spikes from noise events we plotted the manually sorted complex spikes in different colors (Fig 1, light blue – complex spikes and red –noise events). The PC analysis for the first representative neuron achieves good separation in the low-frequency domain, but not in the high-frequency domain (Fig. 1A-C). This is indicated by the high overlap between the complex spike cluster and the noise cluster in the high-frequency projection (Fig. 1C), but not in the low-frequency projection (Fig. 1B). Thus, in some cases, the PC analysis using the low pass filtered signal yielded a better separation of the complex spike cluster. Note that the composite projection (Fig. 1A); i.e., the projection on the first PCs of the low and high frequency data, also clearly differentiated the signal from the noise. For this neuron the separation could be attributed to the low frequency first PC alone, as indicated by the separation of the cluster in the low-frequency axis, but not in the high-frequency axis.

In some cases, the high frequency signal can be informative as well. In contrast to the first neuron, PCA of the second neuron (Fig. 1D-F) provided an example where the complex spikes could be sorted either from the low or high frequency signal. The complex spikes were grouped into an isolated cluster when projecting the data segments on the two leading high frequency principal components (Fig. 1F), or the two leading low frequency principal components (Fig. 1E). The composed PCA achieved good separation as well (Fig. 1D). The cluster of manually sorted spikes (light blue dots in Fig. 1) did not overlap with the cluster of any other event (red dots in Fig. 1). These other events consisted of simple spikes or other non-complex spike events in the extracellular recording. Sometimes these events could be further clustered, as is evident from the example of the small isolated groups in Fig. 1F. These non-CS events were not our primary interest; we therefore denoted them and the corresponding cluster as noise.

In order to compare the low-frequency and high frequency performance, we took data from eight neurons and marked all the complex spikes in the raw data (see Methods) manually. Next, we tested and compared the performances of three forms of PCA: PCA using both low and high frequencies, PCA using low frequency alone and PCA using high frequency alone. The findings showed that using the low-frequency data over high-frequency data for the PC analysis was superior for two reasons: 1. The projection on the low frequency PC yielded better separation of the complex spikes from the noise. 2. The low-frequency data were more effective in detecting the events that enter into the analysis. Below we compare the different PCA methods, followed by an expanded discussion of each low frequency advantage separately.

### Comparison of the three PCA approaches

For each comparison we projected the data on the two relevant PCs and then marked the clusters with a polygon. We then compared this clustering to the manual sorting on the raw data by calculating the recall and precision. The recall is the percentage of complex spikes that were detected in the analysis, whereas the precision is the percentage of the detected events which are indeed complex spikes. The incentive to use precision and recall instead of false negatives and false positives is that the latter is subject to a somewhat arbitrary decision in the threshold setting, which in turn determines the amount of noise (the source of the false positive population) that enters into the analysis. Using false positive and negative measures did not change any of our conclusions.

The recall (Fig. 2A) and the precision (Fig. 2B) were fairly similar for the low frequency and composite measures, and smaller for the high frequency. To compare these analysis results, we calculated the F-measure of each neuron; that is, the harmonic mean of the precision and recall. We compared the performances of each pair of methods and generated a scatterplot of their F-measure for each neuron. Performing the analysis with low-frequency data was significantly better than using the high-frequency data (blue dots in Fig. 2C, F-measure, P=0.0221, permutation test). Similarly, the analysis results for the eight neurons were significantly better when using composed frequencies versus high frequency data (F-measure, P=0.0254, permutation test). As we found for the recall and precision measures separately, the F-measures of the composed and low frequency methods were similar (orange dots in Fig. 2C, F-measure, P=0.4848, permutation test). Note that the composed and low frequency methods only share the same low frequency first PC. This is convincing evidence that the first PC of the low frequencies plays a major role in obtaining the separation of the signal cluster from the noise cluster.

**Figure 2:**
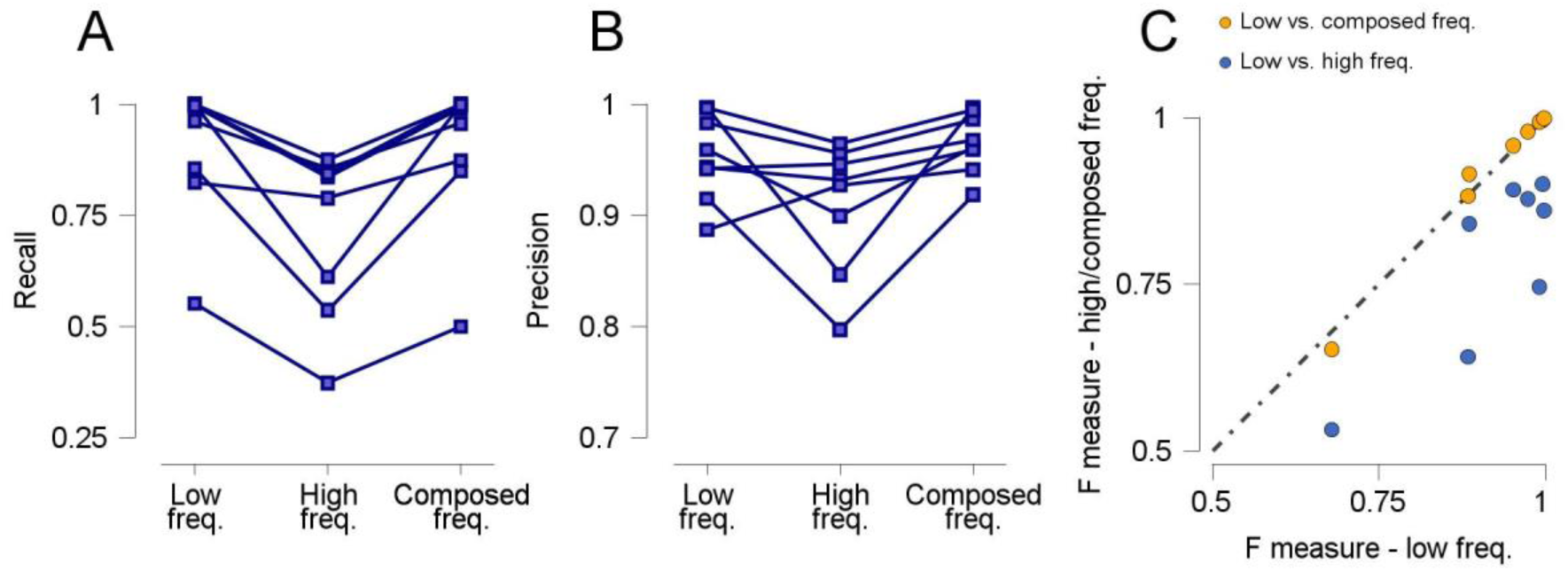
Comparison of three PCA performance. **A.** Percentage of detected events (recall) for three forms of PCA. **B.** Percentage of the detected event made up of complex spikes (precision) for three PCA methods. The lines in **A** and **B** connect the results of the three methods for the same neuron. **C.** Scatter of the F-measure (harmonic mean of recall and precision) for two pairs of methods. Horizontal axis is the F-measure of the low frequency PCA methods. Vertical axis is the F-measure of the composed or high frequency PCA methods. Each dot represents one neuron.

### Advantages of the low frequency signal for clustering complex spikes

To compare the overlap between the histograms of the noise and the complex spikes in the high and low frequency domains, we projected the high frequency segments on the high frequency first PC and the low frequency segments on the low frequency first PC. We then calculated the percentage of complex spikes or noise for each bin, creating histograms of the projection of the signal and the noise on either the high or low frequency first PC (Fig. 3A). The shape of the first PC appears above the histograms of the neuron (for the low frequency and for the high frequency). Note that the high-frequency PC captures the first spike and the spikelets, whereas the low-frequency PC captures the general form of the prolonged large amplitude spike.

**Figure 3:**
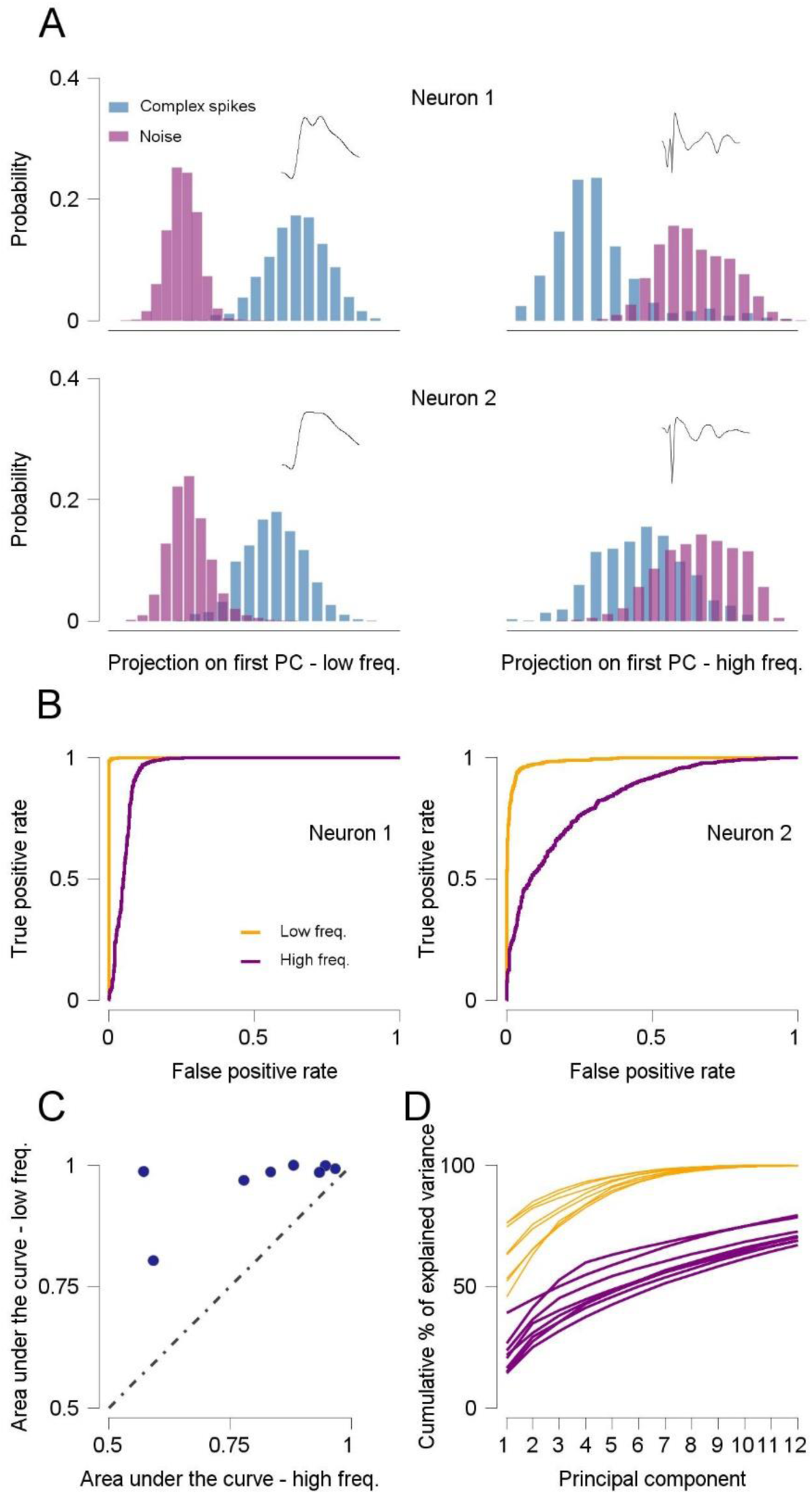
Separating complex spikes from noise using low and high frequency PCs. **A.** Histograms of noise (red) and complex spike (light blue) scores on 1^st^ PC of threshold crossing events. The left and right column show the projection on the low and high frequency 1^st^ PC. The horizontal axis is the probability of events for a given score. Each row is the event projections for different neurons. The shape of the PC the events projected on appears above each graph. **B.** ROC of the high and low frequency data for each neuron in A. Purple and yellow tracers show the analysis for the high and low frequency data. **C.** Scatter of the area under the ROC for all neurons. Each dot represents one neuron and its high (horizontal) and low (vertical) frequency area under the curve. **D.** Scree plot for high and low frequencies. Each line shows the cumulative percentage of the explained variance for one neuron.

The results indicated a better separation of the signal from the noise in the case of the low-frequency PC, as shown by the lesser overlap between the noise and the signal histograms for the low versus the high frequency projections (Fig. 3A left vs. right column). For each neuron, we calculated the receiver operating curve (ROC) for each case (low frequencies or high frequency projections) to quantify the overlap of the histograms (Fig. 3B). For each neuron, we then calculated the area under the ROC for each PCA. The area under the ROC was consistently larger for the low frequencies as indicated in the scatterplot that compares the two measures (Fig. 3C). All dots lie above the equality line, indicating that the projection on the low frequency first PC yields a better separation of the signal histogram from the noise histogram for all tested neurons (Fig. 3C, Wilcoxon test, P=0.0078). Therefore, using the low pass signal for complex spike PC analysis results in a better signal-noise differentiation. A possible explanation is that the first PC of the low frequencies explains a larger fraction of the variance in the data. Generally, the percentage of variance explained by the leading PCs was substantially larger for low versus high frequencies (Fig. 3D). For example, the first PCs explained ~70% of the variance for the low frequencies whereas ~12 PCs were needed to have same amount of explanatory variance for the high frequency. Thus, low frequencies are better at preserving the structure and dispersion of the data, and enable clusters of signal and noise to emerge.

### Advantages of the low frequency signal for detecting complex spikes

The typical first step in sorting spikes consists of setting a threshold and only using events that cross the threshold for further analysis (Lewicki, 1998). We used the low frequency data to set the threshold. We then chose segments in the low-frequency and high-frequency data around these time points of threshold crossing to perform the PC analysis. However, an alternative and common method is to set the threshold on the high-frequency data instead on the low-frequency data. In Figure 4A we depict the advantages of setting the threshold on the low-frequency data. In the high frequency domain, the difference between the amplitude of the signal of the complex spikes and the noise events was small. Often there was no margin to set a threshold that could mainly include complex spikes and exclude simple spikes. Therefore, the threshold on the high-frequency data detects (along with the complex spikes) a considerable amount of noise, such as simple spikes (Fig. 4A, top). In the low-frequency data, the margin between the amplitude of the signal and noise was very large; hence the threshold in the low frequencies will selectively detect complex spikes (Fig. 4A, bottom).

**Figure 4:**
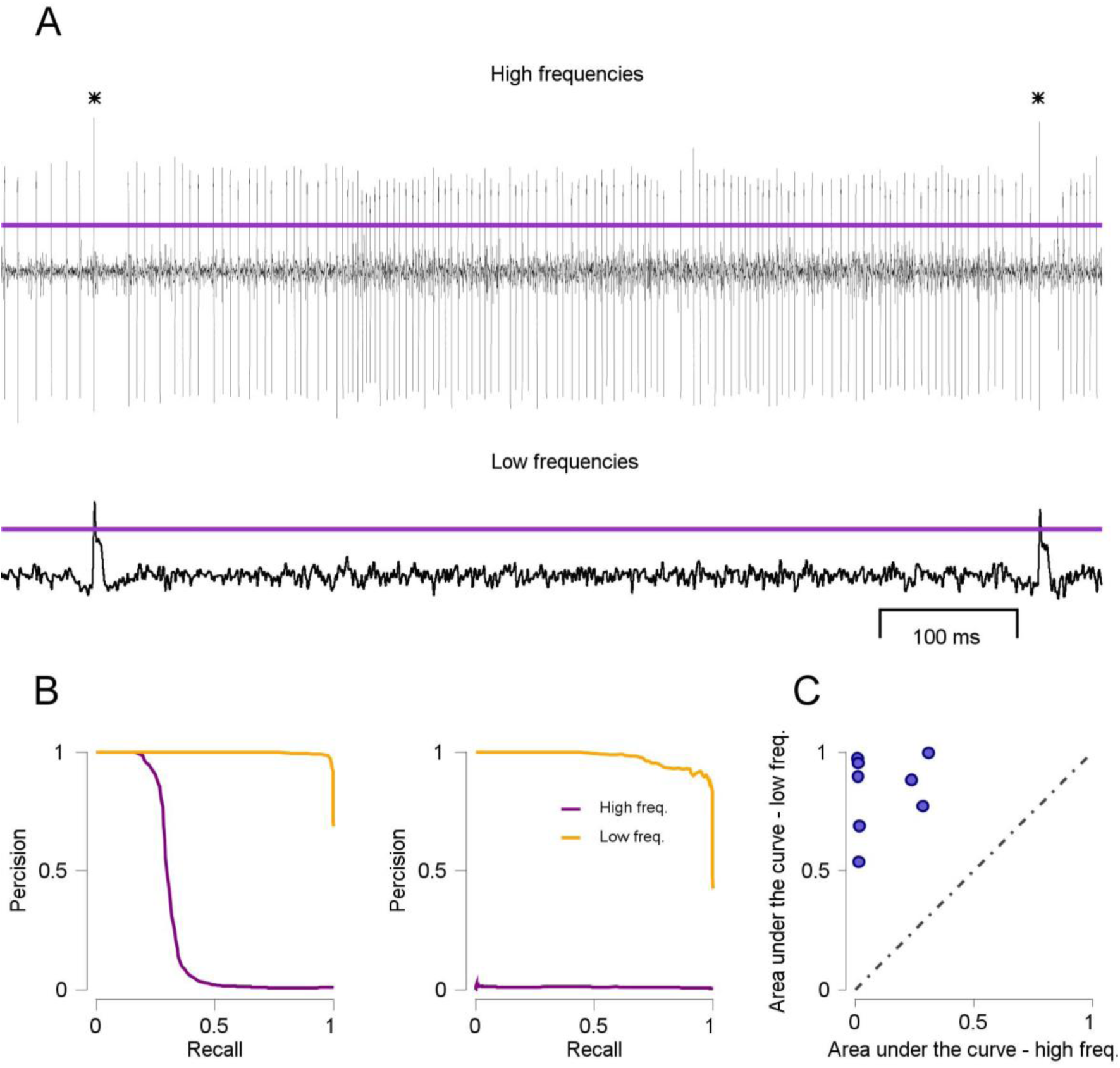
Complex spike detection using low and high frequency data. **A.** Data from a representative neuron in the high (top) and low (bottom) frequency domains. Complex spike events are marked by asterisks. **B.** Precision-recall curves for two neurons. In each graph the recall-precision curves for two threshold settings for the same neuron: threshold on the low frequency data (yellow) and on the high frequency data (purple). **C.** Scatter of the area under the recall-precision curves (AUC), where the horizontal score is the high frequency AUC, and the vertical score is low frequency AUC. Each dot represents one neuron.

To determine which type of filtered data was more effective for setting the threshold, we set possible thresholds on the high and low frequency data of each neuron, and quantified the efficiency of each threshold by examining its recall and precision. Ideally, the threshold should be precise; i.e., it should only detect complex spikes, (precision = ~1) from the high threshold. As we lower the threshold the detection should remain precise while the fraction of detected spikes should increase (i.e. increase in recall). Note that these measures resemble but are not identical to the final recall and precision of the sorting method (Fig. 2) since they precede the clustering phase of sorting.

Figure 4B shows two examples in which the low-frequency data yielded higher precision for each recall, as indicated by the location of the low frequency curve relative to the high frequency curve (Fig. 4B). Both examples show that in the low frequencies there was a range of thresholds in which only complex spikes were detected and almost all the complex spikes were detected. This is indicated by the ability of the detection method to achieve both high precision along with high recall (Fig. 4B orange trace). This was not the case for the high frequencies. For the cell on the left of Fig. 4B for the most extreme thresholds, only complex spikes were detected in the high frequency, as indicated by the large precision value in the left corner. As the threshold is lowered, numerous noise events cross the threshold even before all the complex spikes are detected, as indicated by the decrease in precision as the recall increased. The second example cell (Fig. 4B right) shows a case where there was no threshold that could both include complex spikes without including a substantial amount of noise events, as indicated by the low precision values for all recall values.

To quantify the efficiency of thresholds for each neuron we calculated the area under curve of the recall versus precision plots. The area under the curve was higher for the low-frequency thresholds in all neurons (Fid. 4C, all dots lie above the equality line). Values close to 1 in this analysis indicate that there were thresholds in which all and only complex spikes passed the threshold. Values close to zero indicate that thresholds would necessarily include a few complex spikes with many noise events. The larger the value the more efficient the thresholds that could be used for the neuron. Thus, setting a threshold on the low frequency can detect events with a higher signal to noise ratio.

### Characteristics of low frequency sorted spikes match known properties of complex spikes

To support our claim that the low-frequency events are indeed part of the original complex spike signal (Warnaar et al., 2015) we examined several characteristic features of complex spikes; namely, the similarity between these low-frequency events to well-known complex spike events. In addition to the cells for which we manually verified the complex spike identities, we recorded 46 cells and used a combination of low and high frequency features to detect and sort the complex spikes (see Methods). Complex spikes occur at rate of ~1 Hz (Hobson and McCarley, 1972; Kitazawa et al., 1998; Lang et al., 1999; Mano, 1970). Likewise, the low-frequency events also occur at rate of ~1 Hz (Fig. 5C). Note that these cells have a normal simple spike firing rate of ~80 Hz. In addition, these events were accompanied by a pause in the firing of simple spikes (Fig. 5D), as would be expected for complex spike events (Schonewille et al., 2006; Yang and Lisberger, 2014). We observed a drop in the occurrence of simple spikes for about 15ms after the initiation of the complex spike (time point 0), followed by small increase in the simple spike rate. Note that these pauses were longer than the dead-time of the detection which was 3-6ms. Thus, the low frequency events exhibit the characteristic features of the complex spike signal.

**Figure 5:**
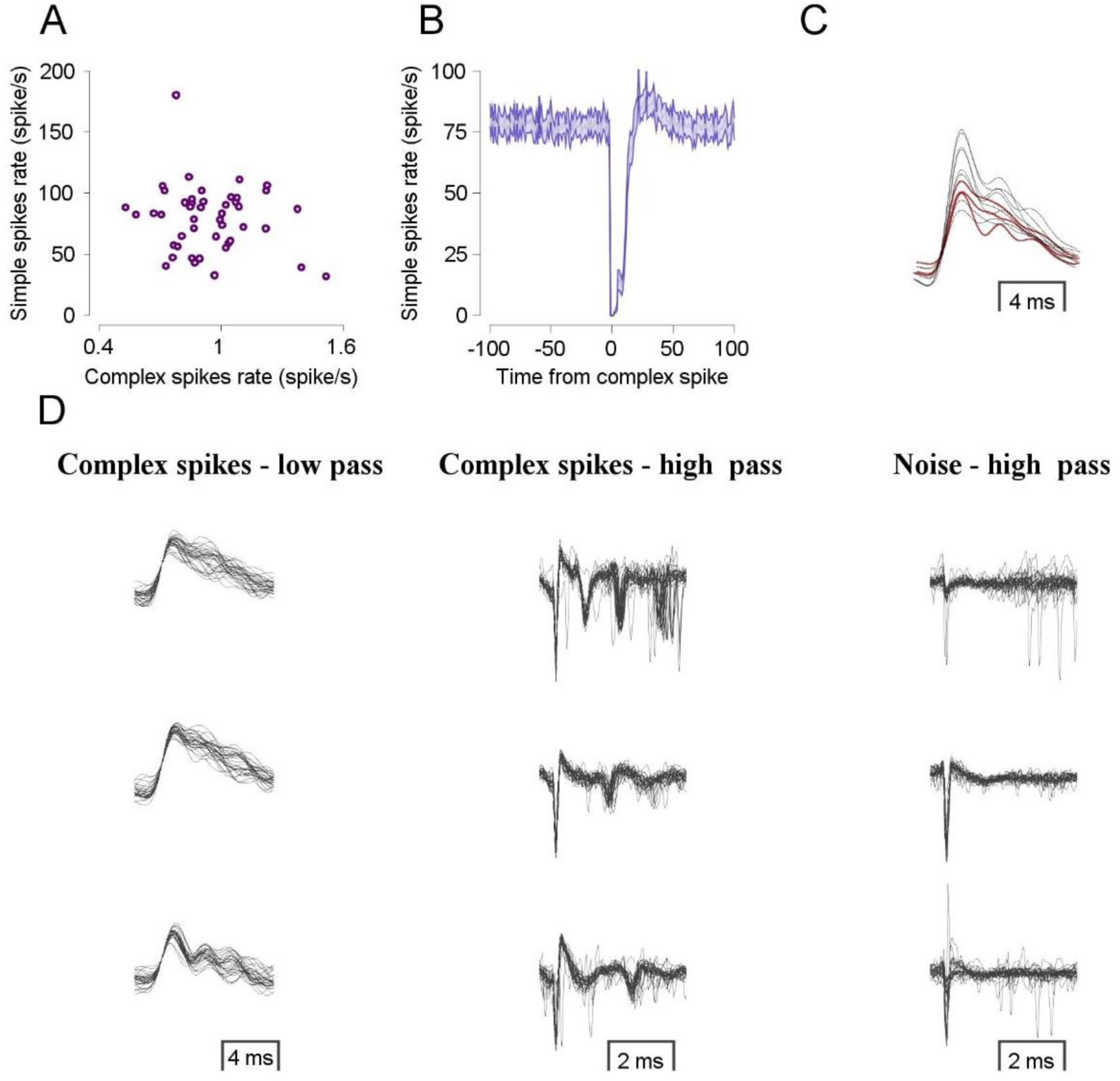
Characteristics of complex spikes measured in the low frequency signal. **A.** Purkinje cells’ spike rate. Each dot represents one cell’s complex spike rate (horizontal) and simple spike rate (vertical). **B.** Simple spike firing rate after a complex spike event (time point zero). **C.** Average of the low-frequency component of the complex spikes. The gray traces show all the neurons that were sorted manually, and red traces show averages from the traces presented in **D**. **D.** Events from three Purkinje cells. Each row represents three type of events (30 events in each subplot) from one cell. The complex spike in the low frequency domain (left column), high frequency domain (middle column), and noise events in high frequency domain (right column).

Next, we searched for neurons in which the high frequency component of the complex spike was clear and distinguishable from the noise (Fig. 5A, right and middle column), and looked at the low-frequency components of these complex spikes (Fig. 5A, left column). The time domain of the 50 randomly selected complex spike events in the low-frequency data was highly self-consistent within the same neuron. Moreover, when comparing the average low-frequency complex spike of these three neurons with the equivalent average complex spike of each one of our eight database neurons, there was a very high similarity in the time domain, in spite of the variability in the amplitude (Fig. 5B). This provides evidence that the low-frequency event of the acquired data, which has a stereotypic shape and time domain, is not the result of some separate source but rather is low frequency signal of the complex spike itself.

## Discussion

The complex spike is a calcium spike (Kitamura and Kano, 2013; Knopfel et al., 1991) generated in the Purkinje cell by input from the climbing fibers (Eccles et al., 1967, 1966). Spikes sorting methods generally rely on high frequency data since spikes are fast events characterized by high frequency. However, because of the highly elaborate dendritic tree of the Purkinje cell, input arriving from the climbing fiber generates a massive large amplitude calcium spike. This makes this type of spike unique in the sense that it can be identified by its low frequency. We exploited this unique feature to improve its sorting. We found that using the low frequency results in better sorting. Further analyses indicated that the using the low frequency leads to more efficient detection and clustering.

### Selecting the preferred analysis procedure

Figure 6 summarizes our analysis procedure. The first step is examination of the complex spikes in the original data to verify whether the complex spikes are composed of a low frequency signal, a high frequency signal, or both. If the only observed frequencies are in the high frequency domain, we recommend setting the threshold and sorting based on the high frequency data. Otherwise, set the threshold on the low frequency data and use the projection of either (our preference) a projection on the 1^st^ PC of the low frequency data and the 1^st^ PC of the high frequency data, or a projection on the 1^st^ PC and 2^nd^ PC of the high frequency data. We have found that in most analyses, the combination marked in red (Fig. 6) was the most effective.

**Figure 6:**
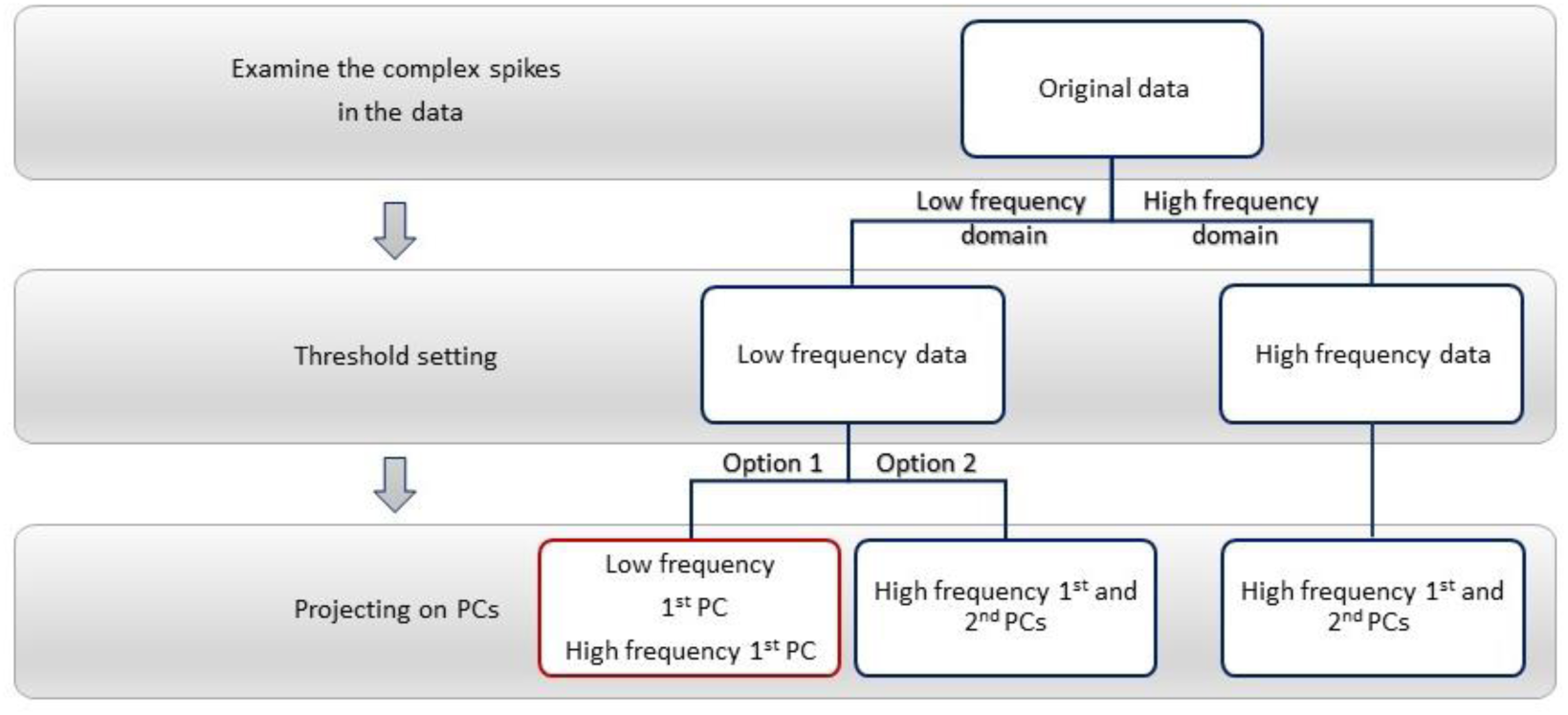
Flowchart summarizing the analysis procedure. Each gray rectangle represents a stage in the analysis procedure. The white rectangles represent the alternative can be used in each stage. Our preference marked in red.

### Physiological aspects of the low frequency signal

The complex spike is characterized by two separate frequency domains where the high frequency domain includes the initial spike and the following spikelets, while these fast events float on a large amplitude event composed of low frequencies. Although these are two separate instances, they correspond to the same comprehensive event in the Purkinje cell known as the complex spike. Thus, we can use either the spikelets (i.e. high frequencies) or the low-frequency spike to detect a complex spike. Our results show that complex spikes are better detected by using the low frequency signal. In some cases, the spikelets are not visible, whereas the large amplitude spike in the low frequency is salient. Hence, in these cases complex spikes can be identified and detected solely by their low frequency signal. Furthermore, multiple complex spikes with noticeable spikelets are challenging to analyze given the variability within spikelets (i.e. the number of spikelets, their timing and etc.). Since our analysis method exploits an additional separate signal of the complex spike (i.e. the low frequency spike) which is not confined to these features, it allows us to record cells with less bias toward complex spikes with highly distinguishable spikelets.

The high and low frequencies are probably qualitatively different and are likely to originate in two different subcomponents of the Purkinje cell (Davie et al., 2008). The high frequencies originate in the proximal axon and propagate to the distal axon and to the soma (Palmer et al., 2010), while the large amplitude low-frequency spike reflects dendritic activity. Computational, anatomical, and functional evidence support the notion that complex spikes encode an error signal needed for motor learning (Albus, 1971; Gilbert and Thach, 1977; Marr, 1969). The complex spike drives long term depression in the parallel fibers to the Purkinje cells (Ito et al., 1982; Suvrathan et al., 2016). In this context the high frequency component of the complex spikes has been used to demonstrate that the complex spikes are not unitary, all-or-none events (Burroughs et al., 2017; Najafi and Medina, 2013). Longer complex spikes with more spikelets induce larger changes in simple spikes and thereby greater changes in behavior (Yang and Lisberger, 2014). Our characterization of the low frequencies of complex spikes provides a way to test whether the low frequencies might also have a functional correlate in awake behaving animals. According to the learning hypothesis, the calcium signals in the dendrites are a critical signal for plasticity. Thus, since the low frequencies are more tightly related to dendritic activity (Davie et al., 2008) they maybe even more strongly linked to learning than the high frequency spike duration. Furthermore, cells in which both high and low frequencies are recorded (Fig. 5) could yield important insights regarding the interplay between the signals in different cell compartments and learning.

### Limitations of using low frequencies for complex spike detection

Using the low frequencies is not always advantageous. In this study we analyzed a subset of randomly chosen Purkinje cells. When we further examined the full set we indeed found cases in which sorting with low frequencies was problematic. In some cases, the data consisted of low frequency noise which made it harder to exploit the low frequencies of the signal, due to the decrease in the signal to noise ratio. For example, the power line frequency (50/60 Hz) falls in the range of the complex spike low frequencies. The local field potentials from the population are also in the same band of the complex spike low frequencies (Pesaran et al., 2018). In these cases, the low frequency signal might still be informative since the shape of the signal may remain different from the noise. Hence, our analysis included the first PC of the high frequency signal, since in these cases, the combination of low and high frequency signals is likely to be rare in the noise but not in the complex spikes.

An additional and infrequent problematic scenario in our dataset occurs when the low frequency signal of the complex spikes is small, resulting in complex spikes characterized by their high frequency domain alone. In this case, when possible, we suggest using the high frequency signal (Fig. 6) alone. A possible factor for the absence of complex spike’s low frequency signal is the position of the electrode. Small changes in the position of the electrode can lead to changes in the complex spikes high (Hensbroek et al., 2014) or low frequencies. In fact, we noticed that in a few cells, small drifts in the electrode position led to loss of the high or low frequencies. These cells thus illustrate cases in which we could be confident that the low frequency signal indeed represented the complex spikes. To overcome this instability, we performed the analysis on small segments of the data, and in each segment we chose the analysis method that was most appropriate for the concurrent signal (Fig. 6). Nevertheless, in the vast majority of cases the low frequency signal was preferable.

Another limitation of the current study is that we compared our method to manual complex spike sorting. Thus, we showed that sorting based on the low frequencies is more consistent with the observer’s judgments. In addition to the authors’ manual clustering an additional observer who was blind to the methods used in this manuscript confirmed our conclusions.

Thus overall we showed that low frequencies can improve spike sorting. This was illustrated for the case of complex spikes in the cerebellum. Low frequencies could also be exploited in other brain regions with large cells that have massive intracellular currents. The characterization of the low frequency signal of complex spikes may contribute to hence revealing the physiological links between complex spikes and behavior.

## Acknowledgements

We thank M. Yarkoni and A. Lixenberg for collecting the dataset. We thank Y. Botschko for technical assistance. Research was supported by the HFSP career development award, the Israel Science Foundation and the European Research Committee.

